# Self-amplifying RNA-based CAR T cell therapy with enhanced duration and multi-genic logic functions

**DOI:** 10.64898/2026.02.18.706661

**Authors:** Yuyang Gu, Jaehoon Choi, Devin Mutha, Christopher Wu, Neil J. Ganem, Mark W. Grinstaff, Wilson W. Wong

## Abstract

Chimeric antigen receptor T (CAR-T) cell therapy is transforming the treatment landscape of hematological malignancies. However, manufacturing with integrating viral vectors is costly, slow, and carries risks including insertional mutagenesis, prolonged B cell aplasia, and other long-term toxicities. Expression of CAR with mRNA can reduce cost, manufacturing timelines, and improve safety. However, the short-lived expression necessitates frequent repeat dosing. Here, we describe a modified self-amplifying RNA (saRNA) platform for engineering CAR T cells with prolonged CAR expression and enhanced durability of tumor control relative to mRNA CAR T cells. In an acute lymphoblastic leukemia (ALL) xenograft model, saRNA CAR T cells achieve superior tumor suppression and prolong survival. Further, a single-strand modified saRNA supports the co-expression of multiple proteins, enabling the construction of advanced CAR systems, such as OR- and AND-gated logic CAR T cells. Together, these results highlight saRNA as a powerful and versatile platform for CAR T cell engineering with favorable safety, efficacy, and accessibility.

## INTRODUCTION

Chimeric Antigen Receptor (CAR) T cell therapy is a transformative immunotherapy with major clinical success, particularly in B-cell derived hematological malignancies. Since the approval of Kymriah® in 2017, seven CAR T therapies have been authorized by the FDA and EMA, five targeting CD19 and two targeting B-cell maturation antigen (BCMA)(*1, 2*), all for oncology. Today, more than 43,000 patients are benefiting from CAR T cell therapies, either commercially or through clinical trials(*3*). Approaches to further advance CAR T cell designs and enhance tumor-targeting specificity and anti-tumor activity include logic CARs or transcription factor overexpression, which require the expression of multiple genes in the same cell(*4-6*). Beyond oncology, CAR T therapies are being investigated for the treatment of autoimmune disorders(*7, 8*), cardiac fibrosis(*9*), HIV(*10*), and senescence-related disease among others(*11, 12*).

Building on these clinical successes, significant opportunities exist to improve CAR T cell performance and patient outcomes. One of the most exciting opportunities is the engineering of CAR T cells using messenger ribonucleic acid (mRNA), which does not integrate into the genome and therefore eliminates the risk of insertional mutagenesis(*13*). Pre-clinical and early clinical results across a range of liquid(*14-16*) and solid(*17-24*) tumors support further development of this transient CAR expression system, which requires less costly infrastructure and a simpler workflow. Furthermore, *in vivo* CAR T cell therapy, in which an intravenously administered mRNA encapsulated within a lipid nanoparticle (LNP) homes to T cells for *in situ* transfection, shows results in some clinical trials against liquid tumors(*25, 26*). This approach further reduces costs, providing rapid treatment without the need for *ex vivo* cell therapy manufacturing, thereby democratizing treatment, reducing healthcare costs, and facilitating compliance for rural or long-distance patients. However, two major hurdles, (1) the durability, especially in the context of complex solid tumor treatments, and (2) the inability to express multiple genes with a single strand, significantly limit mRNA’s potential as the definitive platform for CAR T cell engineering.

The short half-life of mRNA, even when synthesized with modified nucleotides(*27, 28*) (e.g., N1-methylpseudouridine, N1mΨ), necessitates a large dose or multiple doses to be effective, especially in cell therapy context, which increases the risk of adverse side effects, limits global accessibility, and restricts its applications. Other RNA-based platforms, such as circular RNAs (circRNAs) and self-amplifying RNAs (saRNAs), offer substantial promise for increasing protein expression levels and duration(*29*). circRNAs, which exhibit enhanced durability via evading exonuclease degradation, tend to produce lower protein expression levels than mRNA and saRNA(*30*). Moreover, the inefficient circularization reaction and separation challenges hinder circRNA preparation compared to globally-scaled linear mRNA production. Also, circRNA lacking its 5’ cap requires an internal ribosome entry site (IRES) to initiate translation. However, modified nucleotides interfere with IRES RNA folding and function, thus preventing translation of the downstream gene. As such, circRNA is incompatible with modified nucleotides, which prevents its use in applications where RNA containing modified nucleotides is advantageous.

The most common and potent saRNA is derived from the Venezuelan Equine Encephalitis Virus (VEEV), which encodes an RNA-dependent RNA polymerase (RdRp) on the same RNA strand as the sequence for the desired protein cargo. The RdRp produces additional copies of both the full-length saRNA and the subgenomic RNA encoding the gene of interest, thus increasing protein expression levels and duration(*29*). Recently, we and others report that 100% substitution of cytidine with 5-methylcytidine (m5C) in saRNA reduces the innate immune response while supporting replication and enhancing expression levels and duration in various cell types, including primary human immune cells(*29, 31, 32*). Unlike circRNA and mRNA, saRNA efficiently expresses multiple proteins from a single RNA strand with modified nucleotides. This feature enables more advanced cell therapies without the added quality control challenge of using multiple strands, which are increasingly needed for more precise and potent treatments. The extended expression duration, reduced immunogenicity, and polycistronic gene expression capability position m5C saRNA as a powerful and versatile cell and gene therapy platform.

Here, we report a modified m5C saRNA-based CAR T cell platform that achieves prolonged and elevated CAR expression compared with mRNA counterparts. *In vitro*, m5C saRNA CAR T cells induce cell killing for 7 days and secrete higher levels of cytokines than mRNA CAR T cells. In an acute lymphoblastic leukemia (ALL) xenograft mouse model, saRNA CAR T cells achieve complete tumor control over the treatment course while mRNA CAR T cells do not. Leveraging the compatibility between nucleotide-modified saRNA and IRES, advanced CAR logic circuits with OR- and AND-gates achieve the first polycistronic gene expression from a single modified RNA strand in primary human T cells. These saRNA logic CAR T cells execute the intended logic functions in *in vitro* killing assays, illustrating that saRNA readily supports polycistronic expression for sophisticated CAR designs and other complex cell and gene therapies.

## RESULTS

### Self-amplifying RNA enables prolonged and elevated CD19 chimeric antigen receptor expression in primary human T cells compared with mRNA

We first designed CD19 CAR constructs in mRNA and saRNA vectors. For saRNA, the RdRp (VEEV NSP1–4) is upstream of the CD19 CAR, along with an mCherry reporter. (Figure 1A). We produced the modified RNAs by *in vitro* transcription, with N1mΨ in mRNA and m5C in saRNA. We then electroporated RNAs into primary human T cells and quantified surface CD19 CAR expression by flow cytometry over 7 days. mRNA transfected T cells display a rapid rise in CAR expression that peaks at 6 hours, the only time point at which mean fluorescence intensity (MFI) exceeds that of the saRNA group (Figure 1B, F). MFI then declines to baseline by day 2. In contrast, saRNA-transfected T cells reach peak expression at 24 hours and last for 4 days before dropping to baseline (Figure 1B, E). The CAR MFI from saRNA is also significantly higher than that of mRNA from day 1 to day 4. For the percent CD19 CAR^+^ cells, the mRNA group yields higher initial transfection efficiency with a peak on day 1 followed by a rapid decline (Figure 1C, F). The saRNA group starts lower, peaks on day 1, persists for an additional day, and then decreases more slowly (Figure 1C, E).

**Figure 1:**
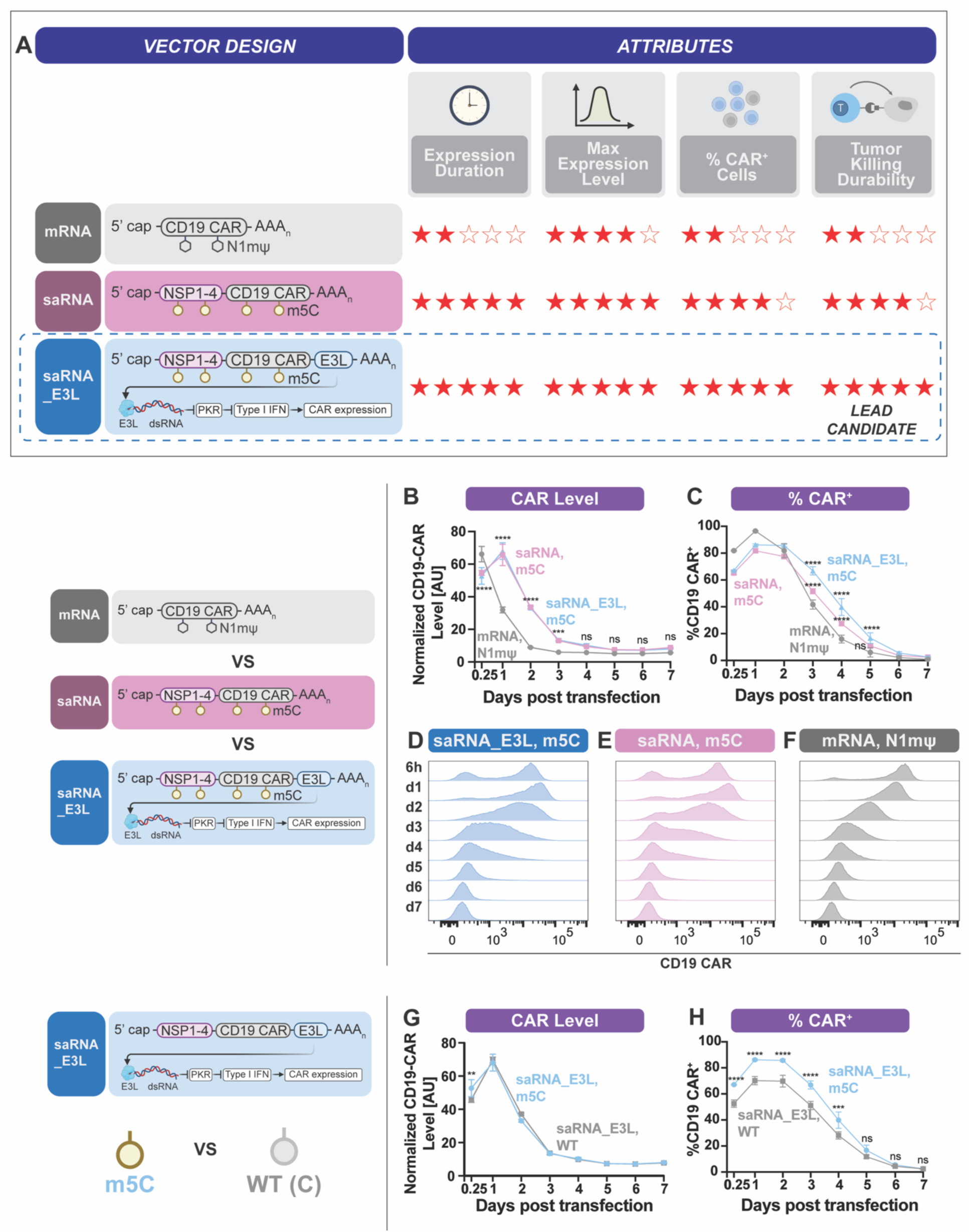
Self-amplifying RNA enables prolonged and elevated CD19 chimeric antigen receptor expression in primary human T cells compared with mRNA. A) Schematic comparing mRNA, saRNA, and an optimized saRNA construct saRNA_E3L for CAR expression duration, maximal expression level, percent CAR^+^ cells over time, and tumor-killing durability. B-C) Time course of CD19 CAR surface expression measured as normalized median fluorescence intensity (MFI) (B) and percent CAR^+^ cells (C) following electroporation with N1mΨ-modified mRNA, m5C-modified saRNA, or m5C-modified saRNA_E3L. CAR was detected by anti-Myc tag staining. MFI was normalized to untransfected cells. Dots and error bars represent mean ± SD (n = 3 biological replicates). Statistical comparisons between m5C-modified saRNA/saRNA _E3L and N1mΨ-modified mRNA were performed by two-way ANOVA with Tukey’s multiple comparisons. * p < 0.05; ** p < 0.01; *** p < 0.001; **** p < 0.0001; ns, not significant. D-F) Representative flow cytometry histograms of CD19 CAR surface expression over 7 days after electroporation with m5C-modified saRNA_E3L (D), m5C-modified saRNA (E), N1mΨ-modified mRNA (F). G-H) Time courses of normalized CD19 CAR MFI (G) and percent CAR^+^ cells (H) after electroporation with wild-type saRNA_E3L or m5C-modified saRNA_E3L. MFI was normalized to untransfected cells. Data are mean ± SD (n = 3 biological replicates). Statistical analysis by two-way ANOVA with Šídák’s multiple comparisons test. Significance notion as in B-C.

To further enhance expression, we co-expressed the vaccinia virus protein E3L, a double-stranded RNA (dsRNA) binding protein that dampens the type I interferon response by limiting protein kinase R (PKR) activation(*33*) (Figure 1A). While the MFI kinetics of the two saRNA constructs overlap closely, the saRNA_E3L construct shows a slower loss of CAR^+^ cells (Figure 1B, C). Histograms for the saRNA_E3L group display a broader peak and a longer high-intensity tail than mRNA (Figure 1D, F). We also tested the incorporation of an additional 3’ UTR upstream of the VEEV 3’ UTR to increase protein expression. Although two β-globin 3’ UTR produces a modest increase in CD19 CAR MFI on day 1, that construct exhibits substantially reduced transfection efficiency compared with the VEEV 3’ UTR only construct (Fig S1A, B). Therefore, inclusion of an extra 3’ UTR tested doesn’t provide an overall benefit. Based on these data, we advanced the saRNA_E3L construct with the VEEV 3’ UTR only for further optimization and functional testing.

Because the CAR includes a C-terminal mCherry, we also tracked mCherry over time. The mCherry MFI from the mRNA group is consistently lower than both saRNA groups, including at 6 hours (Figure S1C-F). The fold difference between saRNA and mRNA groups is larger for mCherry than for surface CAR, likely because mCherry is intracellular, whereas surface display may be limited by membrane trafficking or available surface area.

Next, we evaluated the effect of modified nucleotides on CAR expression. CD19 CAR MFI kinetics for wild-type and m5C-modified saRNA overlap at all time points except at 6 h (Figure 1G). Incorporation of m5C into saRNA increases transfection efficiency and maintains a significantly higher fraction of CAR^+^ cells through day 5 compared with wildtype saRNA (Figure 1H). The m5C modification also reduces IFNβ secretion: IFNβ is below the limit of detection at 6 h for the m5C group versus detectable levels for wild-type saRNA, and remains lower at 24 h, although the difference is not statistically significant (Figure S2A). For mRNA, modification with N1mΨ produces a modest improvement in transfection efficiency and a slightly higher fraction of CD19 CAR^+^ cells on subsequent days, but the effect is less pronounced than for saRNA (Figure S2B). CAR MFI kinetics for wild-type and modified mRNA overlap closely, analogous to the saRNA data (Figure S2C). No IFNβ is detected from N1mΨ mRNA transfected T cells at either 6 h or 24 h, whereas wild-type mRNA elicits measurable IFNβ (Figure S2D). Collectively, the incorporation of m5C in saRNA enhances CAR expression.

### saRNA-based CAR T cells demonstrate more durable cell-killing efficacy *in vitro* than mRNA-based counterparts

With the optimized saRNA construct and nucleotide modification included, we proceeded to functional testing. We performed daily 24-hour co-culture cytotoxicity assays with the B-cell precursor acute lymphoblastic leukemia line, NALM6, for seven days after transfection (Figure 2A). saRNA CAR T cells robustly kill NALM6 cells across the full week, whereas mRNA CAR T cells lose activity within 3 days (Figure 2B). From day 2 post transfection onward, saRNA CAR T cells show significantly higher efficacy than mRNA CAR T cells. In contrast, T cells transfected with mCherry-encoding saRNA and untransfected T cells exhibit minimal killing. Cytokine measurements on day 2 of co-culture reveal significantly higher IFNγ and TNFα secretion from saRNA CAR T cells than from mRNA CAR T at all effector-to-target (E: T) ratios tested (Figure 2C, D). Untransfected T cells produce minimal cytokines. To assess platform versatility, we generated a HER2-targeting saRNA CAR T cell with E3L co-expression and tested it against two HER2^+^ targets, engineered HER2^+^ NALM6 and SKBR3, a human breast cancer cell line. saRNA CAR T cells are cytotoxic in both models across three donors, with minimal basal killing by untransfected T cells (Figure 2E, F).

**Figure 2:**
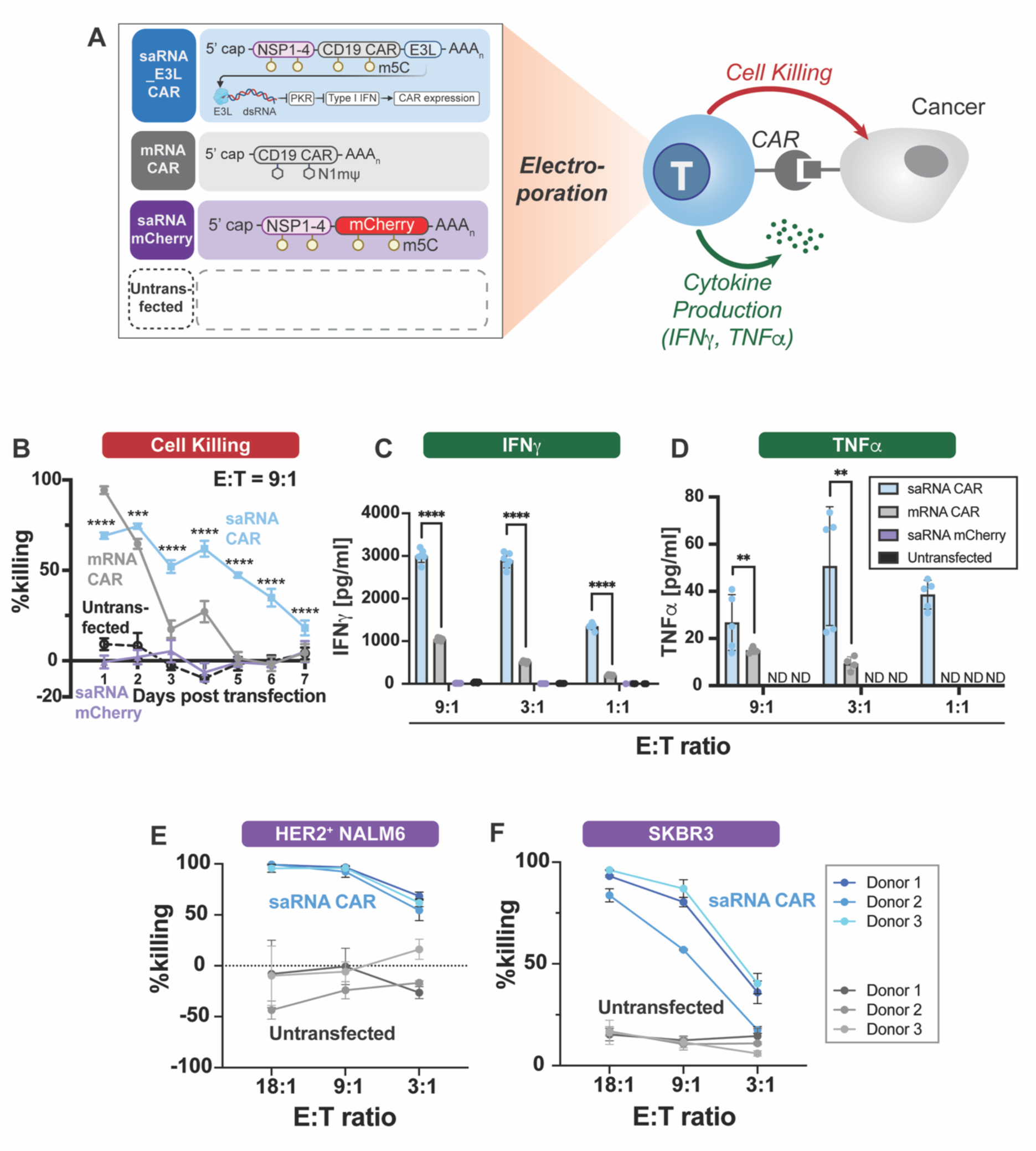
saRNA-based CAR T cells demonstrate more durable cell-killing efficacy *in vitro* than mRNA-based counterparts. A) Schematic of the *in vitro* cell killing assay. Primary human T cells were electroporated with the indicated RNA and cocultured with target cancer cells on specified days post electroporation for 24 h to measure killing. B) Percent killing of NALM6 target cells after 24-hour coculture at effector-to-target (E:T) ratio of 9:1 by mRNA CAR T, saRNA CAR T, saRNA mCherry vehicle-control T cells, or untransfected T cells. Points and error bars are mean ± SD (n = 6 biological replicates). Statistical comparisons versus mRNA CAR T were performed using two-way ANOVA with Tukey’s multiple comparisons. *** p < 0.001; **** p < 0.0001; ns, not significant. C-D) Cytokine concentrations in coculture supernatants on day 2. C) IFNγ at E:T ratios shown; Data are mean ± SD (n = 6 biological replicates). D) TNFα at E:T ratios shown. Data are mean ± SD (n = 5 biological replicates). ND, not detected. Statistical comparisons were performed by two-way ANOVA with Tukey’s multiple comparisons. E-F) Percent killing of HER2^+^ target cells by HER2-targeting saRNA CAR T cells versus untransfected T cells across three donors. (E) engineered HER2^+^ NALM6 cells; (F) SKBR3. Cocultures were performed on day 2 post electroporation and run for 24 h. Points and error bars are mean ± SD (n = 3 biological replicates).

### saRNA-based CAR T cells show superior tumor control versus mRNA-based CAR T cells in an acute lymphoblastic leukemia (ALL) xenograft model

We evaluated *in vivo* efficacy in an established ALL xenograft model(*14*) (Figure 3A, B). We intravenously administered 5×10^5^ luciferase-expressing NALM6 cells into NSG mice. We initiated treatment on day 5 with two weekly injections of CAR T cells or untransfected T cells. We matched the CAR positive dose for both saRNA and mRNA groups (1.5×10^7^, 1×10^7^ cells for two doses, respectively) and used the higher of the total T cell doses for the untransfected T cell dose. We intraperitoneally injected cyclophosphamide 24 h before the second injection(*14*) and monitored tumor burden weekly by IVIS beginning on day 4.

**Figure 3:**
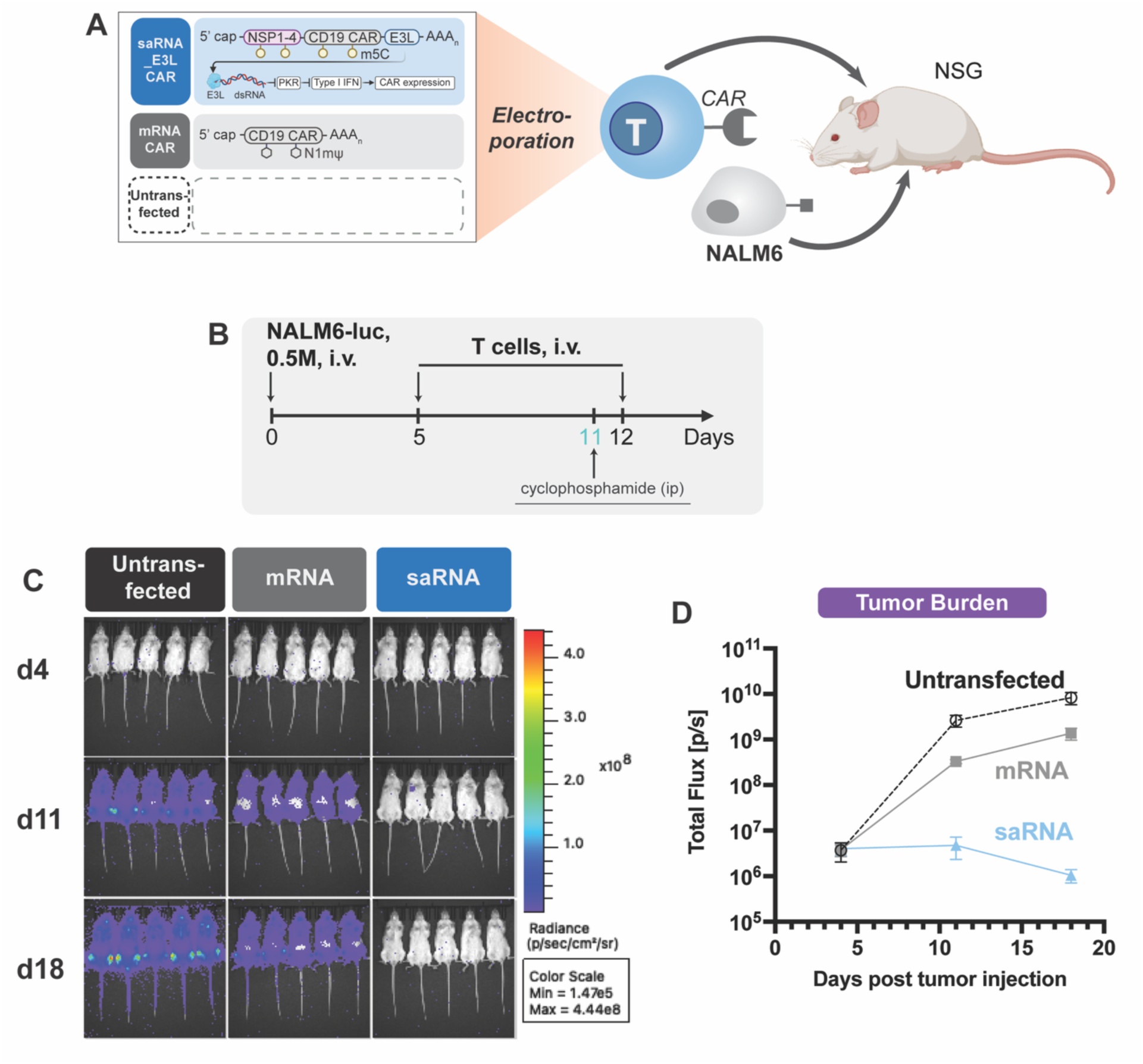
saRNA-based CAR T cells show superior tumor control versus mRNA-based CAR T cells in an acute lymphoblastic leukemia (ALL) xenograft model. A-B) Study schematic. NSG mice received luciferase-expressing NALM6 cell (5×10^5^, i.v.). Mice were randomized and treated with two i.v. doses of CAR T cells or untransfected T cells at the indicated times. Cyclophosphamide was administered i.p. 24 h before the second T cell dose. Tumor burden was monitored weekly by bioluminescence imaging (BLI/IVIS). C) Representative IVIS images on days 4, 11, 18 post tumor injection for untransfected T, mRNA CAR T, and saRNA CAR T groups. D) Whole-animal luminescence (total flux) over time for each group. Data are mean ± SD.

saRNA CAR T cell achieves complete tumor control throughout the treatment course, whereas mRNA CAR T cell provides only partial tumor control with tumors progressing (Figure 3C, D). At lower CAR T cell doses (5×10^6^, 7×10^6^ CAR positive cells, respectively), saRNA CAR T cell again outperforms mRNA CAR T cell in tumor suppression (Figure S3A-C). Within the saRNA arm, CAR T cells manufactured using either Dynabeads or Immunocult activation show comparable efficacy, and both outperform mRNA CAR T cells and untransfected controls in tumor control (Figure S3B-C).

### saRNA achieves polycistronic expression and enables advanced CAR logic circuits

First, we compared IRES, ribosome skipping sequences (P2A, T2A), and subgenomic promoter (SGP) for dual CAR expression (Figure S4A) of CD19 CAR and BCMA CAR. The IRES construct yields the highest double positive population (Figure S4B-H). Considering the structural liabilities of 2A peptides and low expression from SGP, we selected saRNA with IRES for co-expression. We then applied this strategy to build advanced CAR logic circuits targeting ROR1 and HER2. We designed an OR gate and an AND gate (Figure 4C). In the OR gate, each arm contains a complete scFv plus signaling domains. In the AND gate, the anti-ROR1 arm carries only a co-stimulatory domain (CD28) and the anti-HER2 arm carries only the CD3ζ domain, requiring both antigens for full activation. We optimized the saRNA construct order and performed dose titrations to ensure adequate expression (Figure S5A-L). Flow cytometry experiments confirm expression of both OR-gated and AND-gated CARs (Figure 4D, E), and *in vitro* cytotoxicity assays demonstrate the expected logic functions (Figure 4F, G).

**Figure 4:**
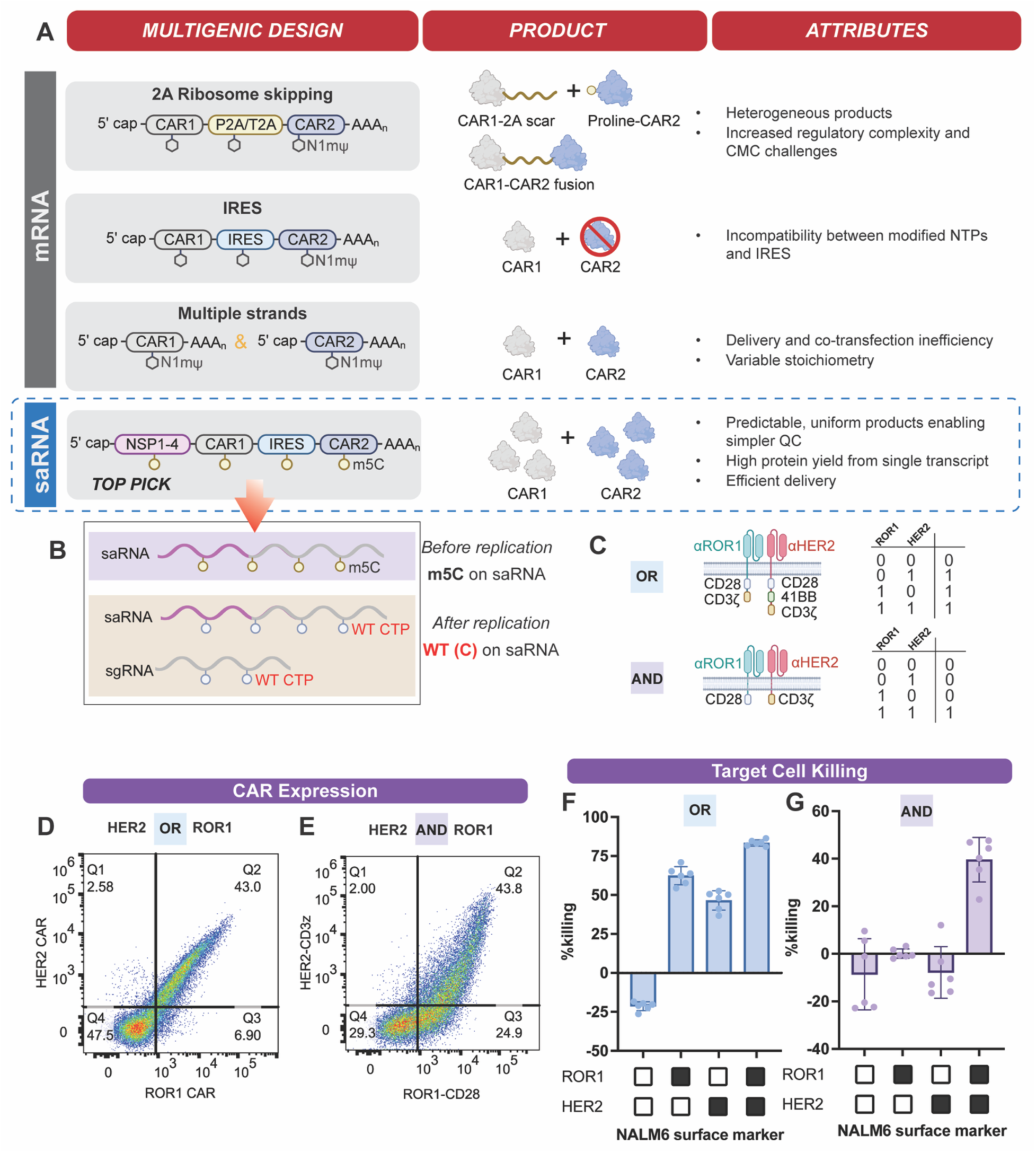
saRNA supports polycistronic expression and enables advanced CAR logic circuits. A) Comparison of strategies for dual CAR expression using conventional mRNA or saRNA vectors, expected protein products, and key attributes. B) Mechanistic schematic illustrating IRES-driven translation compatibility with modified saRNA: during replication, the negative strand saRNA serves as a template for synthesis of the positive strand genomic RNA and the positive strand subgenomic RNA. Subgenomic RNAs are synthesized with wild-type CTP, preserving the native IRES structure, and avoiding disruption by modified NTPs. C) Logic truth table and molecular designs for OR-gated and AND-gated CAR T cells targeting ROR1 and HER2. D-E) Representative flow cytometry plots demonstrating surface CAR expression for OR-gated (D) and AND-gated (E) constructs. F-G) *In vitro* target cell killing by OR-gated (F) and AND-gated (G) CAR T cells at E:T ratio of 9:1. Bars show percent lysis in the presence or absence of cognate antigens. Data are mean ± SD (n = 6 biological replicates).

## DISCUSSION

Linear mRNA-based CAR T cell therapies are advancing across preclinical and clinical settings for oncology(*16, 19-21*) and autoimmune disorders(*34-36*). Early-phase clinical trials with *ex vivo* mRNA produced CAR T cells report favorable safety. Cytokine release syndrome (CRS) was rare: one patient in cited studies experienced grade 1 CRS with fever and arthralgias that resolved within 24 hours without intervention. No neurotoxicity was reported. Clinical responses varied by study and cohort and included complete responses, partial responses, stable disease, and progressive disease, reflecting the advantage of transient mRNA mediated CAR expression. Circular mRNA is an alternative chassis with encouraging preclinical results and longer protein expression(*37-39*), but the manufacturing challenges hinders its progress. Also, reliance on IRES-mediated translation, together with incompatibility between modified nucleotides and IRES(*40*), adds additional complexity, often necessitating separate production of modified and unmodified segments followed by ligation(*41*). Although N6-methyladenosine (m^6^A) dampens the innate immune response, most reported exogenous circular RNA systems use native nucleotides. The tradeoffs with these mRNA technologies include one or more of the following: short-lived activity, higher dose requirement, frequent redosing, increased patient burden, and complex manufacturing. Further, for autologous products, the number of T cells required for multiple infusions can exceed the number that pretreated patients with prolonged cytopenia can provide, and repeated leukapheresis and infusions raise concerns about adherence. Thus, RNA strategies that extend protein expression, sustain CAR T activity, and expand CAR co-expression capabilities are of substantial interest.

Protein expression from saRNA persists significantly longer than mRNA and fills the delivery gap between days with mRNA and months to years with lentiviral/retroviral systems. As a non-genomic integrating technology, saRNA is gaining traction as an alternative RNA platform for preclinical and clinical development. Recent regulatory approvals of wild-type nucleotide saRNA as a COVID-19 vaccine booster in Japan and Europe are a significant milestone(*42, 43*), highlighting the clinical potential of saRNA. m5C saRNA represents a further improvement of the saRNA platform, enabling higher protein expression with reduced interferon signaling *in vitro* and *in vivo* compared to wild-type saRNA and N1mΨ mRNA(*29*).

For the first time, we show that wild-type saRNA and modified m5C saRNA are translationally active in T cells and outperform N1mΨ mRNA. Nucleotide modification increases and sustains CAR positive population (Figure 1H). m5C saRNA CAR T cells maintain robust cell-killing for seven days *in vitro,* whereas N1mΨ mRNA CAR T cells lose substantial activity by day 3 (Figure 2B). *In vivo*, m5C saRNA CAR T cells achieve superior tumor control over the treatment course which shows rapid tumor progression (Figure 3C-D).

Many cell engineering applications require the co-expression of two or more proteins to improve specificity and efficacy. In mRNA constructs, 2A ribosome skipping sequences are commonly used (Figure 4A), but they leave a C-terminal scar on the upstream protein and an N-terminal proline on the downstream protein, which can impair folding and stability(*44-46*). A furin site removes the upstream scar, but 2A cleavage is imperfect(*47-50*). Failed skipping produces fusion proteins that may be dysfunctional. Such heterogeneity complicates Chemistry, Manufacturing, and Controls (CMC) and Quality Control (QC) and increases regulatory complexity. IRES is an alternative, but the downstream cistron cannot be translated in standard N1mΨ mRNA(*40*), effectively eliminating CAR2 and making IRES infeasible in this setting. Delivering separate mRNAs adds co-transfection inefficiency and challenges to control stoichiometry. A single-strand modified saRNA with an IRES overcomes these limitations. During replication of nucleotide-modified saRNA, the RdRp incorporate wild-type nucleotides into RNA species at each step, including negative strand saRNA, positive strand saRNA, and positive strand subgenomic RNA that serves as the template for protein expression (Figure 4B), supporting high level production of both proteins from a single transcript within one cell. Thus, we engineered OR-gated and AND-gated CAR T circuits encoded on a single m5C saRNA strand separated by IRES. *In vitro,* OR-gated CAR T cells demonstrate robust killing in the presence of either or both cognate antigens (Figure 4F), while AND-gated CAR T cells activate only when both antigens are present (Figure 4G), highlighting the saRNA CAR T platform’s potential for multi-targeting efficacy and improved safety by reducing on target off tumor toxicity.

The next logical step is to pair saRNA with advanced delivery systems to enable *in vivo* saRNA-based CAR T therapy and improve on current mRNA-based approaches. In the cited studies(*25, 26*), dosing schedules involved two to three injections per week. Beyond cost and adherence, the feasibility of repeated lipid nanoparticle (LNP) dosing remains uncertain(*51, 52*), and immune responses against LNP components, including anti-PEG antibodies(*53*), have been reported. For *in situ* CAR T generation, logic-gated or drug-inducible CAR designs add an extra safety layer by reducing off-target activity(*4-6*). In autoimmune disease, substantial antigen heterogeneity among patients further favors saRNA’s capacity to support multi-target CARs, while the transient nature allows a prompt immune reset.

In our study, m5C saRNA CAR T cells prolong CAR expression, afford durable tumor killing *in vitro* and *in vivo*, and enable sophisticated CAR T circuits such as OR and AND gates. Collectively, m5C saRNA offers a promising path to improved safety and efficacy for T cell engineering.

## MATERIALS AND METHODS

### Cell lines

NALM6 cells were cultured in RPMI1640 (Corning) supplemented with 10% fetal bovine serum (FBS) (GeminiBio) and 1% Penicillin-Streptomycin (10,000 U/mL) (Gibco). For CD19 CAR T functional *in vitro* killing assays and *in vivo* studies, wild-type NALM6 cells were engineered to express firefly luciferase (Fluc) using a piggyBac system and selected with 100 μg/mL Zeocin™ (Thermo Fisher Scientific). For cell killing assays with HER2 CAR T and HER2/ROR1 logic-gated CAR T, single positive (ROR1^+^HER2^-^, ROR1^-^HER2^+^) and double positive (ROR1^+^HER2^+^) NALM6-Fluc cells were generated by lentiviral transduction of NALM6-Fluc. The HER2 lentivirus encodes the extracellular and transmembrane domains of human HER2. The ROR1 lentivirus encodes the full sequence of human ROR1. Selection was performed with 400 μg/mL Hygromycin B Gold (InvivoGen) for HER2^+^ cells and 2 μg/mL Puromycin (InvivoGen) for ROR1^+^ cells. SKBR3 cells were cultured in McCoy’s 5A medium (Gibco) with 10% FBS and 1% Penicillin-Streptomycin. Cells were maintained at 37 °C in a humidified incubator with 5% CO_2_.

### Plasmid design

mRNA and saRNA vector backbones were adapted as previously reported(*29*) with protein coding sequences replaced by CAR constructs. The CD19 CAR contained an antihuman CD19 single chain variable fragment (scFv) (FMC63 clone), CD28 and CD3ζ signaling domains, and mCherry. E3L sequence reference: NCBI NP_042088.1. The HER2 CAR contained an anti-human HER2 scFv (H3B1 clone), CD28, 41BB and CD3ζ signaling domains, and mCherry. The ROR1 arm of the OR-gate CAR contained an antihuman ROR1 scFv (R12 clone), CD28, CD3ζ, and mCherry. The ROR1 arm of the AND-gate CAR contained an anti-human ROR1 scFv and CD28 only. The HER2 arm of the AND-gate CAR contained an anti-human HER2 scFv, CD3ζ, and mCherry. Unless noted otherwise, all saRNA CAR T cells for functional testing were generated with the saRNA_E3L construct, except for the logic CAR T cells.

### mRNA and saRNA synthesis

Both mRNA and saRNA were synthesized by *in vitro* transcription (IVT). Plasmids were linearized for 16 h at 37 °C using SapI for saRNA and Esp3I for mRNA, then purified with the DNA Clean & Concentrator-25 kit (Zymo Research). IVT was performed with 1 μg of linearized template in a 20 μL reaction using the MEGAscript™ T7 Transcription Kit (Invitrogen™, Thermo Fisher Scientific) per the manufacture’s protocol and incubated at 37 °C for 3 h. Templates were digested with Turbo DNase at 37 °C for 15 min. RNA was purified with the Monarch® Spin RNA Cleanup Kit (New England Biolabs) and eluted in RNase-free water. RNA was stored at -80 °C until use.

### Primary human T cell isolation and culture

Blood was obtained from healthy donors at the Blood Donor Center, Boston Children’s Hospital (Boston, MA). Primary peripheral blood mononuclear cells (PBMCs) were isolated under Boston University Institutional Review Board (IRB) approval. Primary human T cells were enriched from PBMCs using RosetteSep™ Human T Cell Enrichment Cocktail (StemCell) and SepMate™-50 (IVD) according to the manufacturer’s protocol. Cells were cryopreserved in 90% human serum (Innovative Research) and 10% DMSO (Santa Cruz Biotechnology) until use. T cells were cultured in X-VIVO® 15 Serum-free Hematopoietic Cell Medium (Lonza) supplemented with 5% human serum, 1% 1M N-Acetyl-L-cysteine (Sigma), and 0.1% 2-Mercaptoethanol (Life Technologies). For expansion, recombinant human IL-2 was added at 100 U/mL (Hoffmann-La Roche).

### RNA Electroporation

T cells were activated with Dynabeads™ Human T-Activator CD3/CD28 beads (Thermo Fisher Scientific) at 3:1 bead-to-cell ratio. Beads were removed on day 7, and cells were expanded for 2 additional days. On day 9, cells were re-activated with Dynabeads or Immunocult (only if specified) at 0.5:1 bead-to-cell ratio for 24 h, then washed twice and resuspended in Opti-MEM™ (Gibco). Electroporation was performed with 2 μg RNA per 10^6^ cells on a 4D-Nucleofector® X Unit (Lonza) using program EO-115. Cells recovered for 10 min before transfer to complete culture media.

### Flow cytometry

CAR surface expression was assessed by antibody staining. T cells were stained for 30 min at room temperature in FACS buffer (PBS with 2% FBS), protected from light, washed twice and acquired on an Attune NxT Acoustic Focusing Cytometer (Thermo Fisher Scientific). For CAR constructs with myc tag, a rabbit anti-myc monoclonal antibody Pacific Blue™ conjugate (Clone 71D10, Cell Signaling Technology) was used. For CAR constructs with V5 tag, a rabbit anti-V5 Alexa Fluor® 700 conjugate (Clone 1036H, R&D Systems) was used.

### *In vitro* cell-killing assay

Target cells were washed once in PBS and seeded in a 96-well flat bottom plate. CAR T cells were washed once in PBS to remove IL2 and added to targets at the indicated E:T ratios. Cocultures were run for 24 h. Supernatants were collected after centrifugation at 350 g for 5 min and stored at -80 °C for cytokine analysis. Cell viability and percent killing was quantified using Bright-Glo™ luciferase assay kit (Promega).

### Cytokine release assay

Cytokines were quantified by enzyme linked immunosorbent assay (ELISA) using Human IFNγ ELISA MAX™ Standard Set (BioLegend), Human TNFα ELISA MAX™ Standard Set (BioLegend), and Human IFNβ ELISA MAX™ Standard Set (BioLegend), following the manufacturers’ protocols. Colorimetric absorbance was measured on a SpectraMax® iD3 plate reader (Molecular Devices).

### Mouse xenograft studies

All procedures were approved by the Boston University Institutional Animal Care and Use Committee (IACUC) (PROTO201800600). NSG mice (5-6 weeks old, Jackson Laboratory) received 5×10^5^ luciferase-expressing NALM6 cells i.v.. Tumor burden was monitored by bioluminescence imaging (BLI/IVIS) weekly beginning on day 4. Treatment with CAR T cells or untransfected T cells began on day 5 and was administered weekly. Cyclophosphamide (60 mg/kg) was given i.p. 24 h before the second T cell injection. For IVIS, mice received D-luciferin (15 mg/mL, 200 μL, i.p., Gold Biotechnology) and were imaged 10 min later.

### Statistical Analysis

Statistics were performed in GraphPad Prism v10.6.1. Data are presented as mean ± SD. Statistical tests and significance notation are reported in the figure legends.

## ACKNOWLEDGEMENTS

This work was supported by the National Institutes of Health [grants R01CA296810 (WW, MWG), U01CA265713 (WW, MWG), R01EB029483 (WW), and R01EB038005 (WW, MWG)], National Science Foundation Trailblazer Award (MWG), National Science Foundation Graduate Fellowship (CW), Boston University Kilachand Award, and BU IVIS Imaging Core. We thank Joshua McGee for the critical reading of the manuscript.

## CONTRIBUTIONS

YG, JC, MWG, and WWW conceptualized the study. YG designed and optimized the RNA constructs and performed *in vitro* testing, with input from JC. YG and JC designed the *in vivo* studies. YG prepared tumor cells and CAR T cells for the *in vivo* studies. JC conducted IVIS imaging and intraperitoneal injections for the study in Figure S3. DM performed all injections and IVIS imaging for Figure 3 and the intravenous injections for Figure S3. CW assisted with data analysis. YG drafted the manuscript, and all authors reviewed and edited the final version. MWG and WWW secured funding. MWG, WWW and NJG supervised the study.

