## Supplemental Information for "Self-amplifying RNA-based CAR T cell therapy with enhanced duration and multi-genic logic functions"

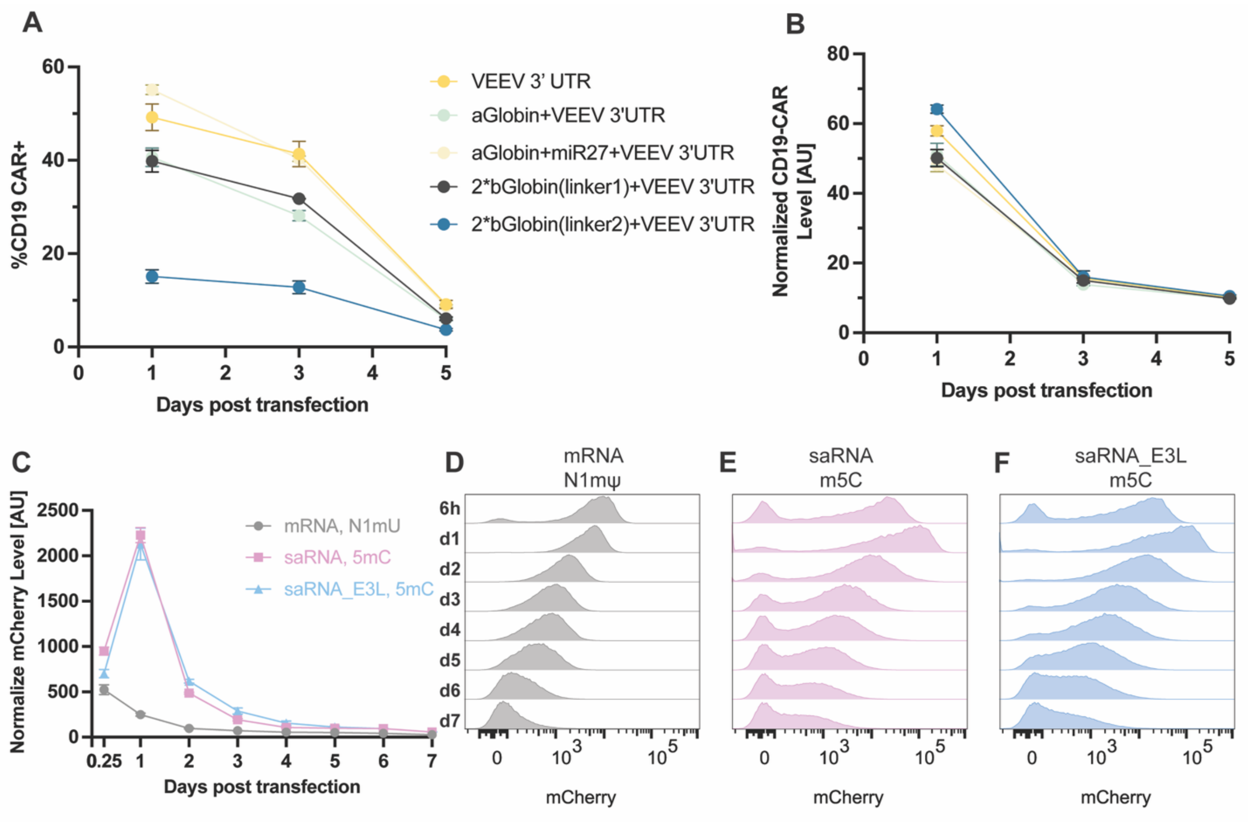


**Supplementary Figure 1**. **A)** Time course of %CAR$^+$ cells after electroporation with saRNA_E3L constructs bearing the indicated 3' UTR inserted upstream of the VEEV 3' UTR. **B)** Time course of normalized CD19 CAR MFI for the same constructs in (A). **C)** Time course of normalized mCherry median fluorescence intensity (MFI) in primary human T cells after electroporation with N1mΨ-modified mRNA, 5mC-modified saRNA, or 5mC-modified saRNA_E3L. mCherry is fused to the C terminus of the CD19 CAR. MFI was normalized to untransfected cells. Points and error bars represent mean ± SD (n = 3 biological replicates). **D-F)** Representative flow cytometry histograms of mCherry over 7 days after electroporation with N1mΨ-modified mRNA (D), 5mC-modified saRNA (E), or 5mC-modified saRNA_E3L (F).

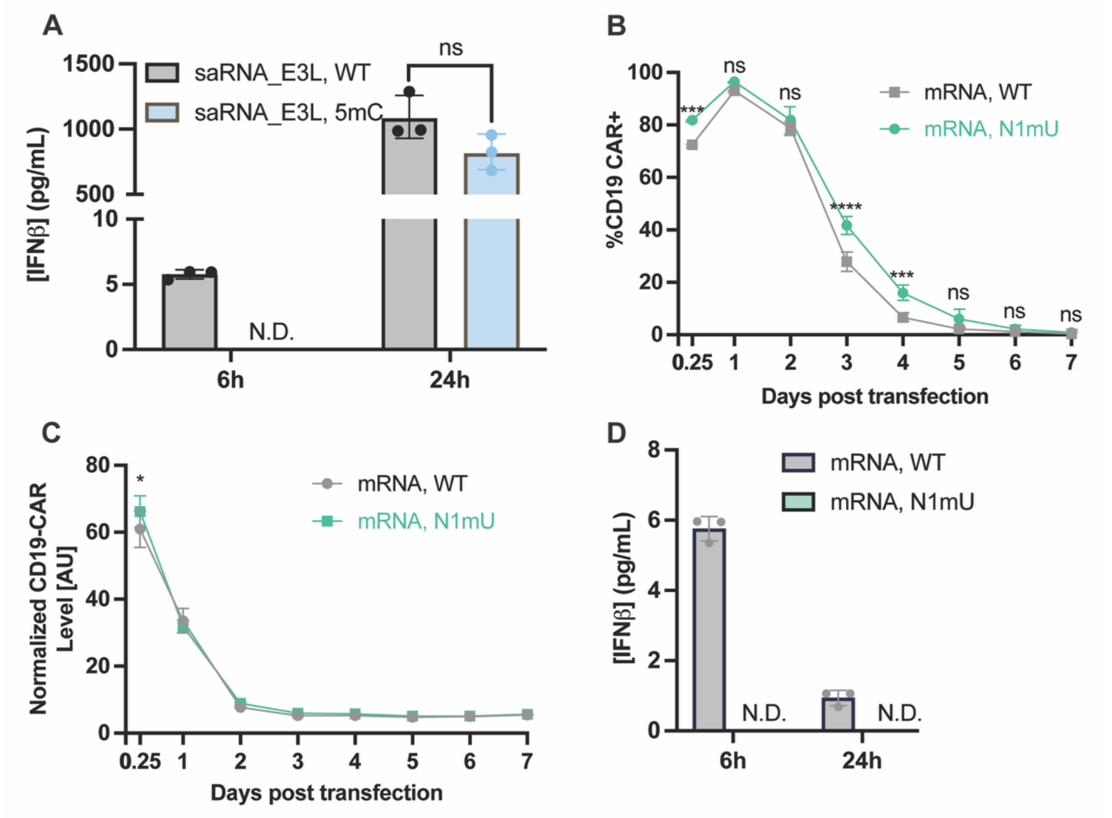


**Supplementary Figure 2. A)** IFNβ secretion from primary human T cells at 6 h and 24 h after electroporation with wild-type saRNA_E3L or 5mC-modified saRNA_E3L. Data are mean ± SD (n = 3 biological replicates). Statistical analysis by two-way ANOVA with Tukey's multiple comparisons correction. ns, not significant. **B-C)** Time courses of %CAR$^+$ cells (B) and normalized CD19 CAR MFI (C) after electroporation with wild-type mRNA or N1mΨ-modified mRNA. MFI was normalized to untransfected cells. Data are mean ± SD (n = 3 biological replicates). Statistical analysis by two-way ANOVA with Šídák's multiple comparisons test. * $p < 0.05$; *** $p < 0.001$; **** $p < 0.0001$; ns, not significant. **D)** IFNβ secretion from primary human T cells at 6 h and 24 h after electroporation with wild-type mRNA or N1mΨ-modified mRNA. ND, not detected.

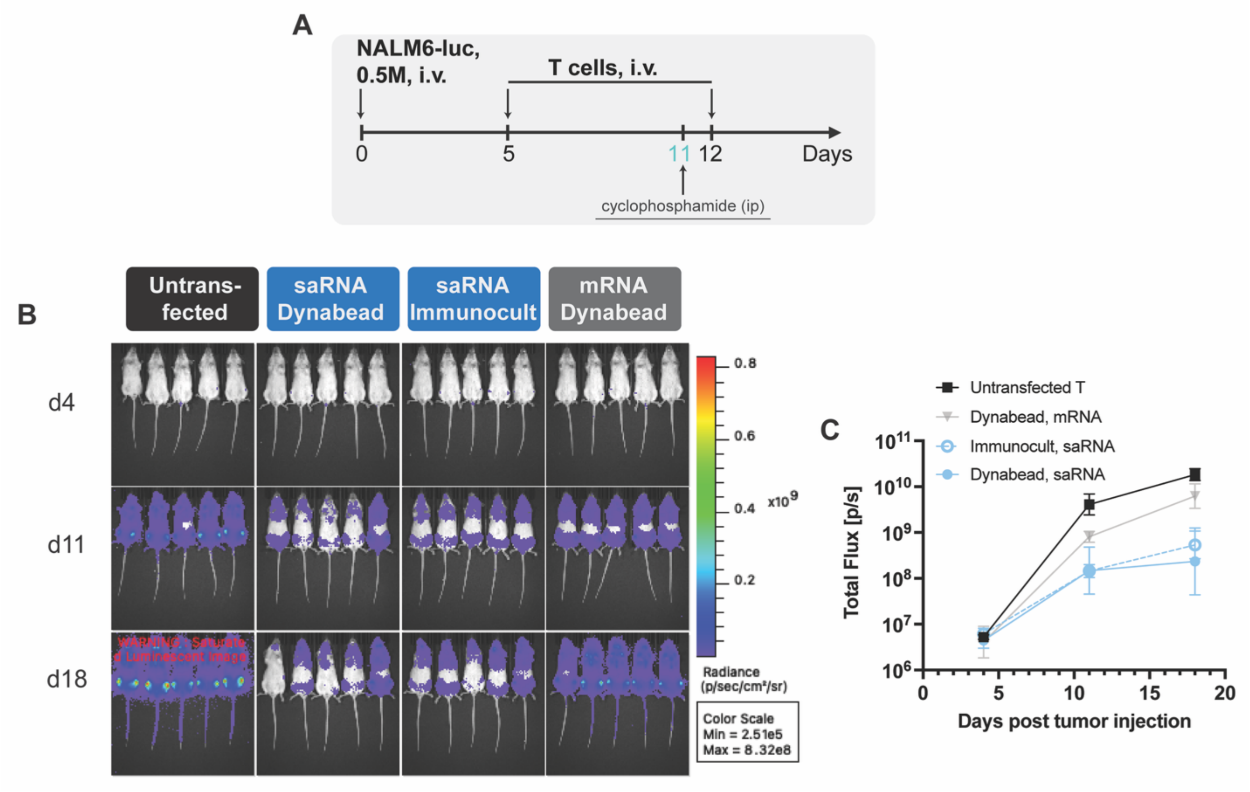


**Supplementary Figure 3. A)** Study schematic. NSG mice received luciferase-expressing NALM6 cell ($5{\times}10^5$, i.v.). Mice were randomized and treated with two i.v. doses of CAR T cells or untransfected T cells at the indicated times. Cyclophosphamide (60 mg/kg) was administered i.p. 24 h before the second T cell dose. Tumor burden was monitored weekly by bioluminescence imaging (BLI/IVIS). **B)** Representative IVIS images on days 4, 11, and 18 for untransfected T, mRNA CAR T (Dynabeads re-activated), and saRNA CAR T manufactured with Dynabeads or Immunocult re-activation. **C)** Whole-animal luminescence (total flux) over time for each group. Data are mean ± SD (n = 5 per group).

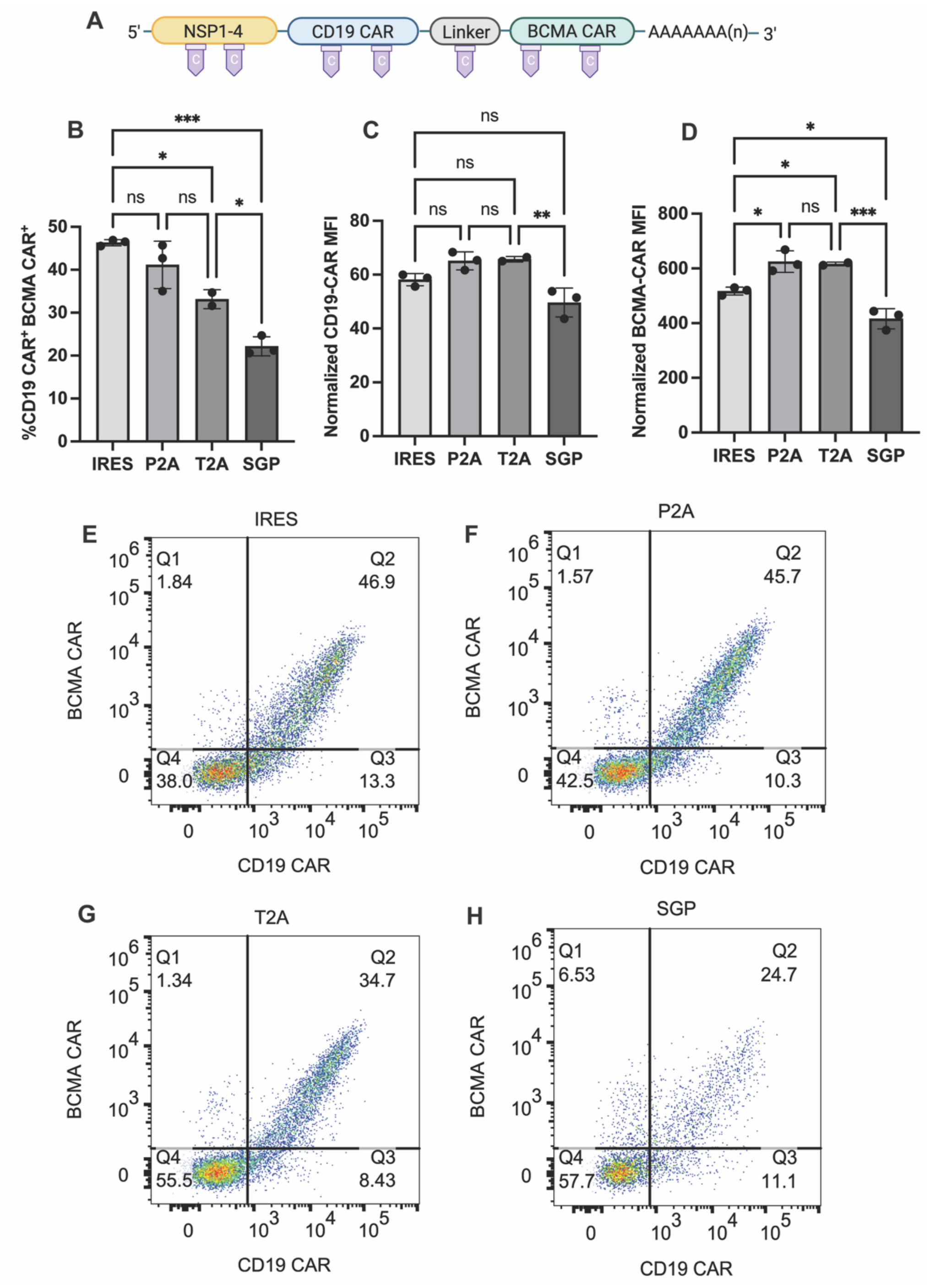


**Supplementary Figure 4. A)** Schematic of dual CAR constructs using different linkers. **B-D)** Double positive frequency for CD19 CAR and BCMA CAR (B), normalized CD19 CAR MFI (C), and normalized BCMA CAR MFI (D) on day 1 after electroporation with 5mC-modified saRNA. MFI values were normalized to untransfected cells. Points and error bars show mean ± SD. Statistical comparisons were performed by one-way ANOVA with Tukey's multiple comparisons correction. * $p < 0.05$; ** $p < 0.01$; *** $p < 0.001$; ns, not significant. **E-H)** Representative flow cytometry plots of BCMA and CD19 CAR expression using IRES (E), P2A (F), T2A (G), or SGP (H).

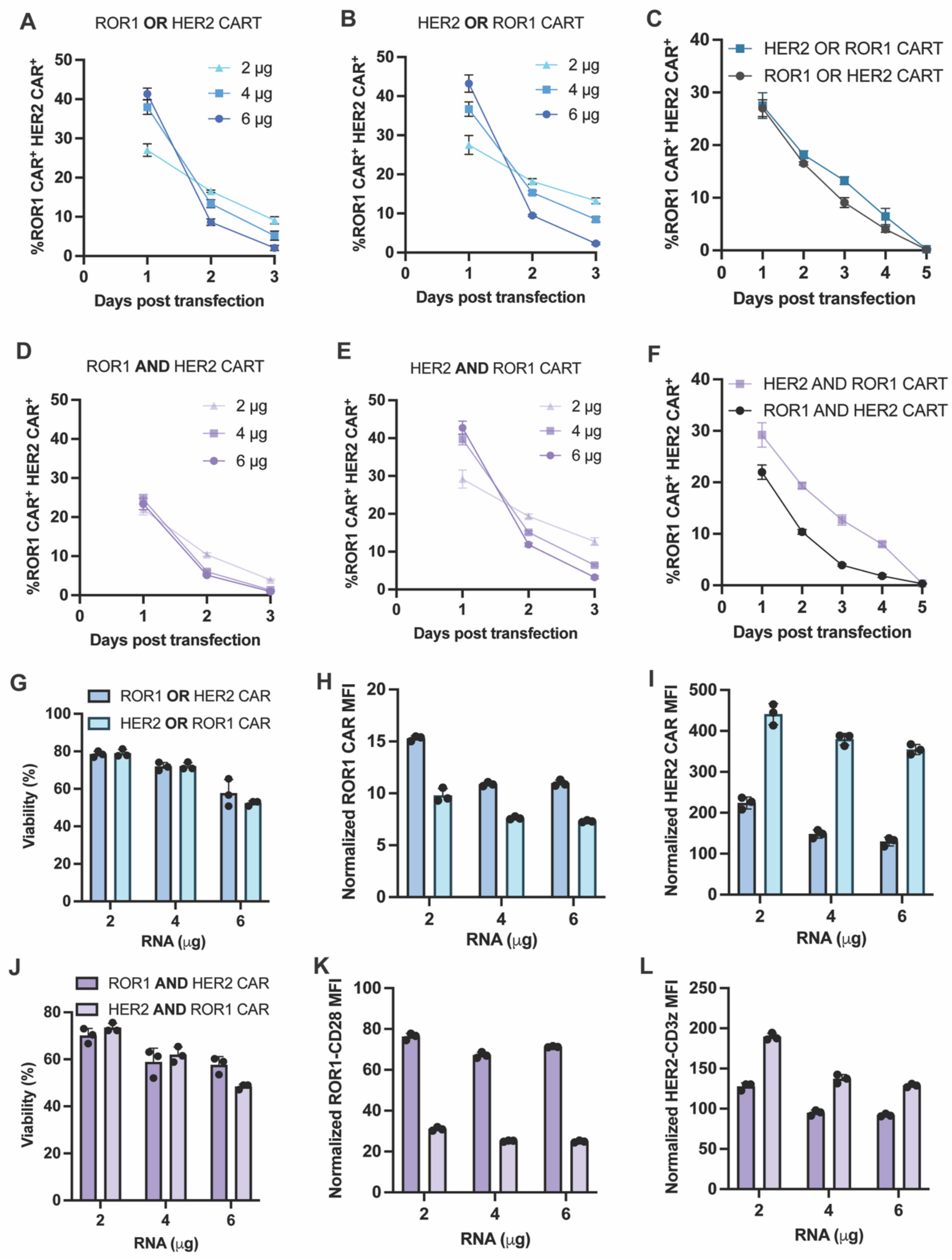


**Supplementary Figure 5. A-C)** Time courses of the double positive fraction co-expressing both CARs for OR-gated constructs after electroporation with 5mC saRNA at the indicated doses, comparing the two arm orders. Dose for (C) is 2 μg per $10^6$ cells. **D-F)** Time courses of double positive fraction for AND-gated constructs at the indicated dose, comparing arm order. Dose for (F) is 2 μg per $10^6$ cells. **G-I)** Viability (G), normalized ROR1 CAR MFI (H), and normalized HER2 CAR MFI (I) for OR-gated constructs across doses and arm orders on day 1 post transfection. **J-L)** Viability (J), normalized ROR1 CAR MFI (K), and normalized HER2 CAR MFI (L) for AND-gated constructs across doses and arm orders on day 1 post transfection. MFI values were normalized to untransfected cells. Points and error bars denote mean ± SD (n = 3 biological replicates).
